# Rapid and sensitive single cell RNA sequencing with SHERRY2

**DOI:** 10.1101/2021.12.25.474161

**Authors:** Lin Di, Bo Liu, Yuzhu Lyu, Shihui Zhao, Yuhong Pang, Chen Zhang, Jianbin Wang, Hai Qi, Jie Shen, Yanyi Huang

## Abstract

Prevalent single cell transcriptomic profiling (scRNA-seq) mechods are mainly based on synthesis and enrichment of full-length double-stranded complementary DNA. These approaches are challenging to generate accurate quantification of transcripts when their abundance is low or their full-length amplifications are difficult. Based on our previous finding that Tn5 transposase can directly cut-and-tag DNA/RNA hetero-duplexes, we present SHERRY2, a specifically optimized protocol for scRNA-seq without second strand cDNA synthesis. SHERRY2 is free of pre-amplification and eliminates the sequence-dependent bias. In comparison with other widely-used scRNA-seq methods, SHERRY2 exhibits significantly higher sensitivity and accuracy even for single nuclei. Besides, SHERRY2 is simple and robust, and can be easily scaled up to high-throughput experiments. When testing single lymphocytes and neuron nuclei, SHERRY2 not only obtained accurate countings of transcription factors and long non-coding RNAs, but also provided bias-free results that enriched genes in specific cellular components or functions, which outperformed other protocols. With a few thousand cells sequenced by SHERRY2, we confirmed expression and dynamics of *Myc* in different cell types of germinal centers, which were previously only revealed by gene-specific amplification methods. SHERRY2 is able to provide high sensitivity, high accuracy, and high throughput for those applications that require high number of genes identified in each cell. It can reveal the subtle transcriptomic difference between cells and facilitate important biological discoveries.

## Background

Many experimental methods for transcriptome profiling by next generation sequencing (RNA-seq) have been developed to cover various scales of input samples, ranging from bulk samples [1, 2] to single cells [3–5] or even subcellular components [6, 7]. High quality single-cell RNA-seq (scRNA-seq) data can be used to reveal the kinetic details of gene expression and transitions between cell states or types [8–10]. Prevalent scRNA-seq methods mainly rely on template switching and pre-amplification of complementary DNA (cDNA). However, large-scale scRNA-seq techniques, commonly operated in micro-droplets or wells, have relatively low sensitivity [11]. Single-tube based scRNA-seq approaches can typically produce higher coverage for low-abundance genes, but they still suffer from quantification bias due to insufficient reverse transcription and GC imbalance during amplification. Besides, their complex experimental methods are generally unsuitable for large-scale studies.

We have reported a highly reproducible and rapid library preparation method for RNA-seq, SHERRY, which can be applied to minute amount of RNA samples [12]. The development of SHERRY was based on the recent discovery that Tn5 transposase can bind and cut RNA/DNA hetero-duplexes directly. With slight modifications, SHERRY could also be applied to various clinical metatranscriptome applications, such as identification of SARS-CoV-2 and other pathogens [13].

Although SHERRY was applied to process single cells and achieved less biased quantification of gene expression in comparison with other scRNA-seq methods, the results still exhibited clear coverage bias toward the 3’ -ends of transcripts, relatively low sensitivity, and low tolerance to endogenous DNA. In this work, we present an optimized method, SHERRY2, which addresses the limitations of SHERRY and is fully compatible with single cells and single nuclei with low RNA content. In comparison with prevalent RNA-seq methods, SHERRY2 showed higher sensitivity, better concordance with reference data, greater reproducibility between replicates, and superior scalability, allowing the method to be used to process a few thousand single cells per batch and thus reducing the time required to conduct experiments.

## Results

### SHERRY2 provides high sensitivity and even coverage across gene bodies for scRNA-seq

For scRNA-seq, RNA degradation and incompleteness of reverse transcription (RT) are two major factors that reduce gene detection sensitivity and coverage evenness. Although adding random RT primers facilitates the coverage of long transcripts, it requires removal of ribosomal RNA, which is incompatible with scRNA-seq [13]. Spiking template-switching oligonucleotides also provides more uniform coverage, but this strategy has limited detection sensitivity and specificity [12].

We altered various experimental parameters of the original SHERRY protocol for both bulk (**Additional file 1: Fig. S1-2, Additional file 2-3**) and single cell inputs (**Additional file 1: Fig. S3A**). To protect RNA from degradation, we lowered the concentration of free Mg^2+^, either by reducing the amount of total Mg^2+^ or adding more dNTP to chelate Mg^2+^ ions [14], and observed significant improvement of the coverage evenness of RNA-seq. To facilitate cDNA synthesis, we screened different reverse transcriptases and found that SuperScript IV (SSIV), working at a relatively high temperature with a low Mg^2+^ concentration, could better overcome the secondary structure of RNA and hence simultaneously enhanced the sensitivity and uniformity of RNA-seq.

When RNA-seq was conducted using pictogram-level RNA inputs, sufficient amount of Tn5 transposome was important for high sensitivity, and Bst 3.0 DNA polymerase filled the gap left by Tn5 tagmentation more effectively than other enzymes. The protocol was insensitive to many experimental conditions, including the usage of single strand DNA binding proteins [15], Tn5 inactivation, the concentration of extension polymerase, and the usage of hot-start polymerase.

We named this optimized protocol SHERRY2. Using RNA extracted from HEK293T cells as input, we compared the performance of SHERRY2 and the original SHERRY protocol. At the 10-ng level, both protocols identified more than 11,000 genes at saturation. At the 100-pg level, SHERRY2 performed better than SHERRY and detected 5.0% more genes at 0.6-million reads (**Additional file 1: Fig. S2A**). In addition, SHERRY2 greatly diminished 3’-end coverage bias (**Additional file 1: Fig. S2B**) and increased the unique mapping rate for 10-ng and 100-pg inputs (**Additional file 1: Fig. S2C**). We also constructed a bias-free RNA-seq library using 200-ng total RNA input via the conventional fragmentation-and-ligation method with the NEBNext E7770 kit (NEBNext). For 100-pg input, the gene overlap between NEBNext and SHERRY2 was greater than that between NEBNext and SHERRY (81.7% vs 78.4%) (**Additional file 1: Fig. S2D**), and the gene expression results of NEBNext and SHERRY2 were also more closely correlated (R=0.70 vs R=0.65) (**Additional file 1: Fig. S2E**).

The SHERRY2 protocol for scRNA-seq contains only four steps: reverse transcription, Tn5 tagmentation, gap-filling through extension, and PCR amplification. The entire SHERRY2 protocol can be completed within 3 hours, one hour less than the original SHERRY protocol, and still held its competence in costs (**Additional file 1: Fig. S3B**). Other high-sensitivity scRNA-seq methods such as SmartSeq2 may require much more time and more steps to be completed [3] (**Fig. 1A**). The one-tube workflow of SHERRY2 is readily scalable to high-throughput applications. SHERRY2 was able to detect 10,024 genes (FPKM >1) on average within a single HEK293T cell at 1-million reads. When subsampling to 0.2-million reads, SHERRY2 still detected 8,504 genes on average, which was 1,622 (23.6%) more than SHERRY and 886 (11.6%) more than SmartSeq2 (**Fig. 1B**). In addition, the reproducibility of SHERRY2 was significantly higher than that of SHERRY or SmartSeq2 (**Fig. 1C**) due to its simplified workflow and stable performance. Moreover, the evenness of gene body coverage for SHERRY2 was much higher than that of the original SHERRY protocol (0.84 vs 0.72) and was comparable to that of SmartSeq2 (0.84) (**Fig. 1D**). The exonic rate of SHERRY2 was also improved in comparison with that of SHERRY, likely due to the higher RT efficiency of the newly developed method (**Fig. 1E**).

**Figure 1.**
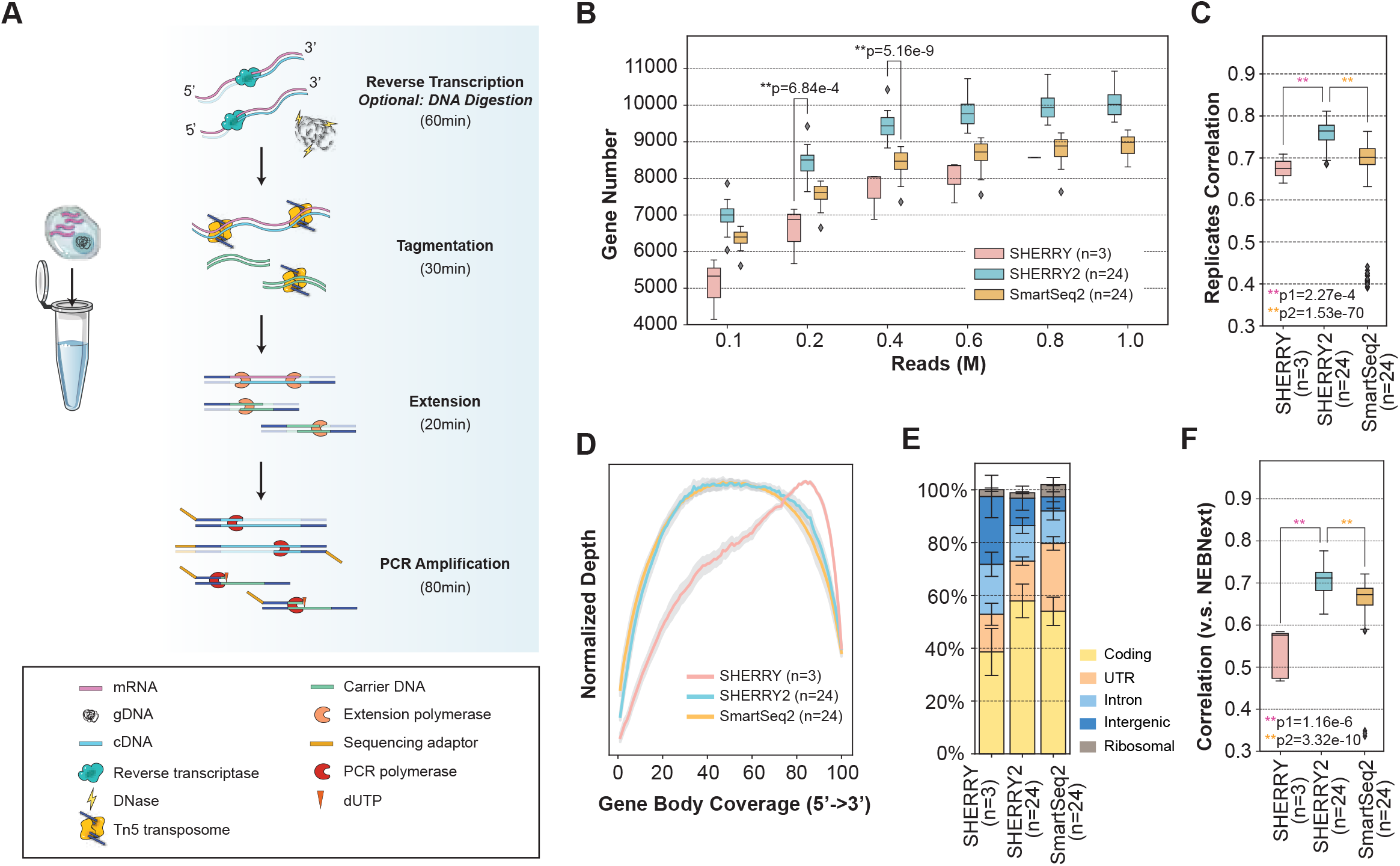
The workflow and general performance of SHERRY2 on single cell RNA-seq. (A) The workflow of SHERRY2 for scRNA-seq. Poly(A) tailed RNA is firstly released from single cells and reverse transcribed. The resulting RNA/cDNA hetero-duplex is then tagmented by Tn5 transposome, followed by gap-repair and indexed PCR. Optionally, chromatin can be digested during lysis. The entire protocol is performed in one tube and takes 3 hours. (B) Gene number (FPKM>1) with SmartSeq2, SHERRY2 and SHERRY when subsampling reads to 0.1, 0.2, 0.4, 0.6, 0.8 and 1 million reads. (C) Pairwise correlation of gene expression within replicates for the three scRNA-seq protocols. The correlation R-value was calculated by a linear fitting model with normalized counts of overlapped genes. (D Gene body coverage detected by the three scRNA-seq protocols. The gray region represents the standard deviation of the normalized depth among replicates. (E) Components of reads that were mapped to different regions of the genome using the three scRNA-seq protocols. The error bars show the standard deviation. (F) Gene expression correlation between single HEK293T cells and 200-ng RNA extracted from HEK293T cells. Single-cell data were acquired by the three scRNA-seq protocols. Bulk RNA results were acquired by the standard NEBNext protocol. The correlation R-value was calculated by a linear fitting model with normalized gene counts. The samples in (B-F) are single HEK293T cells. The p-values in (B, C, F) were calculated by the Mann-Whitney-U test.

Last but not least, scRNA-seq with SHERRY2 exhibited superior accuracy, as demonstrated by the significantly higher correlation between the SHERRY2 gene expression results and NEBNext libraries in comparison with that of SmartSeq2 (R=0.71 vs R=0.67) (**Fig. 1F**), since NEBNext fragmented mRNA before cDNA synthesis and amplified cDNA with very limited cycles which theoretically resulted in negligible bias at transcriptome level. Especially, SHERRY2 showed high tolerance to GC content and was insensitive to the length of transcripts (**Additional file 1: Fig. S4**). Unlike SmartSeq2, for which the gene overlap and expression correlation with bulk RNA-seq showed clear declines when GC-content was greater than 40%, SHERRY2 maintained these parameters at high and constant levels (82.6% overlap and R=0.76) until the GC content reached 60%. Transcript length did not influence the accuracy of SHERRY2 or SmartSeq2, although SmartSeq2 exhibited a small degree of intolerance for transcripts longer than 800 bases.

### scRNA-seq for low RNA-content cells

For low RNA-content cells, such as immune cells [16], we found that removal of intergenic DNA contaminations by DNase treatment was especially crucial for SHERRY2 scRNA-seq. In such cells, the open DNA regions of disassembled chromatin might be favored over RNA/DNA hybrids during Tn5 tagmentation. When DNase was omitted from the SHERRY2 protocol, more than 50% of reads sequenced from single mouse lymphocytes (**Additional file 1: Fig. S5A**) were mapped to intergenic regions, and only around 10% of reads were exonic reads (**Fig. 2A**).

**Figure 2.**
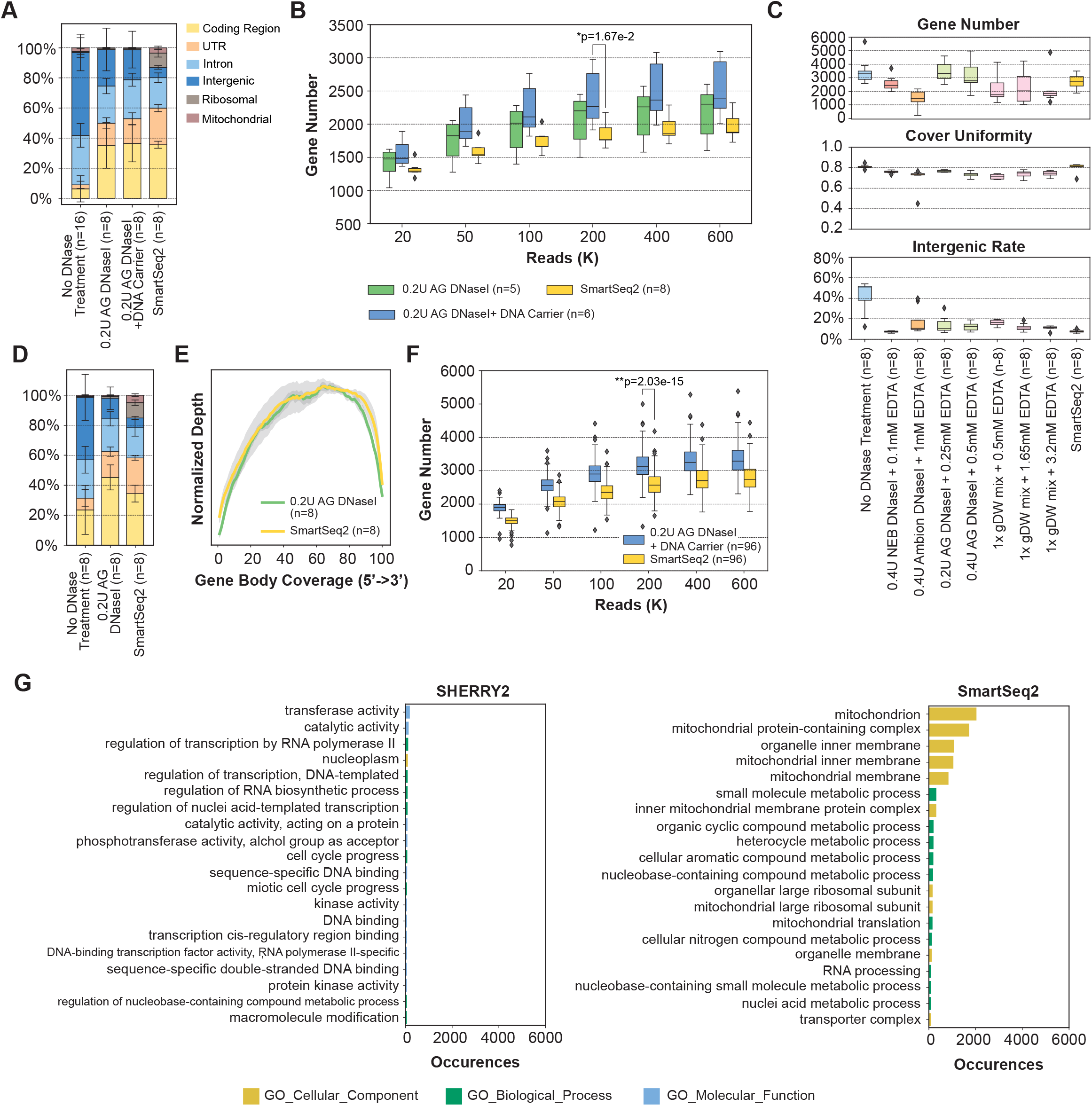
scRNA-seq of low RNA-content samples with SHERRY2. (A) Proportions of genome regions covered by reads from SHERRY2 without DNase treatment, SHERRY2 with AG DNase I addition, SHERRY2 with AG DNase I and DNA carrier addition, and SmartSeq2. (B) Gene number (FPKM>1) detected by SHERRY2 with AG DNase I addition, SHERRY2 with AG DNase I and DNA carrier addition, and SmartSeq2 when subsampling to 20, 50, 100, 200, 400, and 600 thousand reads. Only samples with intergenic rate lower than 25% were counted. Samples in (A, B) were single lymphocyte cells from murine eyeball blood. (C) Library quality of SHERRY2 tested with different DNases, including gene number (FPKM>1) at 0.25-million reads, coverage uniformity across gene body and percentage of reads that were mapped to intergenic regions. The labels below the figure indicate the amounts and names of the DNases, as well as the EDTA concentration that was added during DNase inactivation. SmartSeq2 was also performed as a reference. (D) Components of reads covering different genome regions detected by SHERRY2 without DNase treatment, SHERRY2 with optimized AG DNase I, and SmartSeq2. (E) Gene body coverage detected by SHERRY2 (with AG DNase I) and SmartSeq2. The gray region shows the standard deviation of the normalized depth among replicates. (F) Gene number (FPKM>1) detected by SHERRY2 (with AG DNase I and DNA carrier) and SmartSeq2 when subsampling to 20, 50, 100, 200, 400, and 600 thousand reads. (G) Gene ontology analysis of genes that only detected by SHERRY2 (left) or SmartSeq2 (right). The top 20 most commonly occurred GO terms were shown. Samples in (C-G) were single B cells isolated from murine GC light zones. The p-values in (B, F) were calculated by the Mann-Whitney-U test. The error bars in (A, D) show the standard deviation.

Different DNases performed differently in SHERRY2 scRNA-seq. We tested five DNases (**Additional file 1: Fig. S6A**) and found three (NEB, Ambion, and TURBO DNase I) that worked and inactivated at higher temperatures increased the intergenic rate unexpectedly, and this effect was probably due to RNA degradation at high temperatures with excess Mg^2+^ in the reaction buffer. In contrast, AG DNase I and gDW mix, which worked at room temperature, yielded ideal results.

We confirmed that all the five DNases could digest more than 99.5% of DNA (30-ng) by simply utilizing divalent ions of their respective storage buffer (**Additional file 1: Fig. S6B, Additional file 4**). Without adding extra divalent ions, the intergenic rates of single germinal center (GC) B cells for all DNases were less than 20% (**Fig. 2C**). Among the DNases, AG DNase I retained high sensitivity for gene detection, and more than 60% of reads were mapped to exon regions (**Fig. 2D**), while the evenness of coverage was not affected (**Fig. 2E**).

Next, dU-containing carrier DNA, which could not be amplified by dUTP-intolerant polymerase, was added to further improve the efficiency of tagmentation of RNA/DNA hybrids. With carrier DNA, SHERRY2 detected 3,200 genes at saturation (0.6-million reads) for single GC B cells (**Fig. 2F**), and the number of detectable genes increased from 2,301 to 2,393 on average for single lymphocytes, with an exonic ratio comparable to that of SmartSeq2 (**Fig. 2A, 2B**). Moreover, we examined the genes that were only detected by one method for single GC B cells, and found that SmartSeq2 was preferential to capture genes participated in mitochondrial function (**Fig. 2G**). Based on these results, chromatin digestion and the addition of carrier DNA were included in the standard SHERRY2 protocol and the step of chromatin digestion would consume another 20 minutes.

### Selection dynamics in germinal centers profiled by SHERRY2

SHERRY2 can be easily scaled to thousands of single cells per batch, owing to its simplified procedure. The GC is a transient structure that supports antibody affinity maturation in response to T cell-dependent antigens, and it contains diverse cell types with complex dynamics. Histologically, the GC can be separated into two micro-compartments, the dark zone and the light zone [17, 18]. By surface phenotyping, cells in the two compartments can be distinguished through *CXCR4, CD83* and *CD86* markers [19–21], with light-zone cells being *CXCR4^lo^CD83^+^CD86^+^* while dark-zone cells *CXCR4*^+^*CD83*^lo^*CD88*^lo^. GC cells cycle between the dark zone and light zone states. Dark zone cells are highly proliferative and undergo somatic hypermutation, which generates a range of affinities against antigens. In the light zone, these B cells compete with each other for survival factors and help signals, which are mainly derived from follicular helper T cells. Those B cells that have acquired higher-affinity B cell receptors are selected to differentiate into plasma cells (PC) or memory B cells (MBC) or cycle back to the dark zone [18, 22–24]. Recently, gray zone, consisting of *CXCR4*^+^*CD83*^+^ cells with distinct gene expression patterns, was discovered and found to be involved in GC recycling [25]. The complex spatiotemporal dynamics of the GC and their underlying mechanisms are incompletely understood. To this end, sensitive scRNA-seq methods that can be used to detect gene expression with less bias are highly desirable.

We profiled 1,248 sorted *CXCR4*^lo^*CD86*^hi^ GC light zone cells with SHERRY2, and 1,231 (98.6%) high-quality cells were retained for downstream analysis (**Additional file 1: Fig. S5B**). The gene expression levels of *Cd19, Ccnd3, Fas, Cd86* and *Cxcr4* were consistent with flow cytometry gating (**Additional file 1: Fig. S7A**), and no batch effect was observed (**Additional file 1: Fig. S7B**).

Unsupervised clustering identified seven clusters (**Fig. 3A**), two of which belonged to the gray zone, which was defined by co-expression of *Cxcr4* and *Cd83*, as well as the on-going cell division (enriched *Ccnb1*) [25] (**Fig. 3B**). We observed the expected down-regulation of *Bcl6* and *S1pr2*, the signature genes of GC B cells [26, 27], in memory B cell precursors (MPs) and plasma cell precursors (PPs). Specifically, *Ccr6* was exclusively enriched in MPs [28], while *Irf4* was up-regulated in PPs, which was known to be mediated by NF-κB pathway downstream of *Cd40* stimulation [24]. It’s worth noting that our results exhibited such *Cd40* signaling effects as well (**Additional file 1: Fig. S7C**). Besides, *Icam1* and *Slam1* which were reported to be activated by *Cd40* [29] were also observed (**Additional file 1: Fig. S7D, Additional file 5**). The relatively low expression levels of *Prdm1* (not shown) and *Gpr183* in PPs were consistent with the early stage of plasma cell development. In total, 1,071 genes significantly up- or down-regulated in specific clusters were identified.

**Figure 3.**
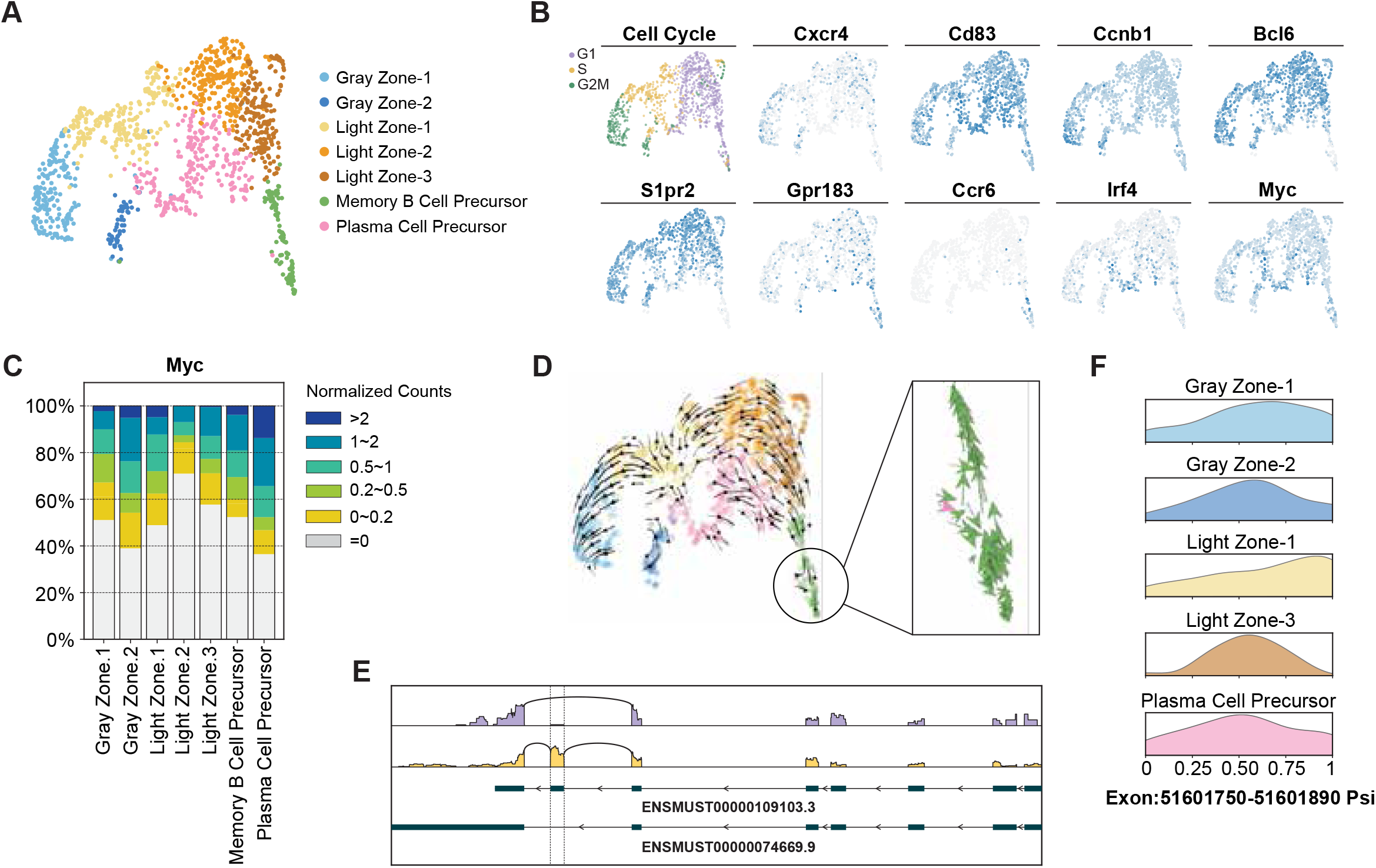
Mouse germinal center profiled by scRNA-seq through SHERRY2. (A) Clustering of single B cells from murine GC light zones visualized by UMAP plot. The library was prepared by SHERRY2 (with AG DNase I and DNA carrier). Different colors indicate distinct cell types. (B) Cell cycle and marker gene expression of different cell types marked on a UMAP plot. The gradient colors correspond to the normalized counts of a specific gene ranging from 0 (white) to 1 (blue). (C) Distribution of *Myc* gene expression in different cell types. Different colors indicate different intervals of normalized *Myc* counts. The percentages of cells within the clusters falling into corresponding intervals were counted. (D) Dynamic process of the GC light zone indicated by vector fields of RNA velocity on a UMAP plot. The expanded region shows the velocity vector of each cell. The colors correspond to the same cell types as annotated in (A). (E) Isoforms of the *Hnrnpab* gene. The top two lines show isoforms from two example cells that rarely and preferentially used the highlighted exon in *Hnrnpab* transcripts. The bottom two lines show the isoform structures of *Hnrnpab* transcripts that include or exclude the exon. (F) Inclusion ratio distribution of the highlighted exon in (E) in different cell types. Only cell types represented by more than 10 cells after filtering are shown.

The high sensitivity of SHERRY2 enabled detection of *Myc* in 588 (47.8%) single GC light zone B cells. Using fluorescent protein reporting, *Myc* was found to mark light-zone cells destined for dark zone re-entry [30], although *Myc* expression *per se* had been difficult to identify in specific cell types by low-sensitivity scRNA-seq approaches [31]. Consistent with previous findings [25, 29], *Myc* expression was significantly higher in PPs (**Fig. 3B, Additional file 1: Fig. S7E**) and active in the gray zone cells (**Fig. 3C**). Light Zone-1 also had a relatively higher portion of *Myc*^+^ cells, which are probably those destined for cyclic re-entry to the dark zone [30]. MPs also contained some cells that expressed *Myc*.

RNA velocity analysis (**Fig. 3D**) suggested that Light Zone-1 contained cells selected for dark zone re-entry, which were migrating to the gray zone and had *Myc* expression characterized by burst kinetics (**Additional file 1: Fig. S7F**). In addition, cells that appeared to have just entered the light zone were also identified. A few velocity vectors that moved to MPs were mixed in PPs, and these vectors were in the same direction with the down-regulation of Myc. According to the velocity analysis, the aforementioned *Myc*-expressing MPs seemed to have a tendency to cycle back to the GC, suggesting that some MPs with *Myc* up-regulation have the potential to re-participate in GC reactions.

We then assembled the BCR sequence for each cell to screen the usage of Igh variable sequences, which were assigned in 1,101 (89.4%) cells. As expected [32], IGHV1-72 dominated the NP-reactive GC response, and the coupled light chain was mainly IgL rather than IgK (**Additional file 1: Fig. S8A, S8B**). In addition, we identified CDR1 and CDR2 regions in 269 (24.4%) and 493 (44.8%) cells in which Igh variable sequences were assigned, respectively (**Additional file 1: Fig. S8C**).

SHERRY2 revealed differences in the usage frequencies of exons across cell types. The usage of a particular exon (chr11: 51,601,750-51,601,890) within the *Hnrnpab* transcript (**Fig. 3E**), which is widely expressed and encodes a protein that mainly functions in processing pre-mRNAs, was significantly biased among GC clusters. As shown in **Fig. 3F**, Light Zone-1 cells favored inclusion of this exon.

### Superior performance of SHERRY2 applied in snRNA-seq

Single nucleus RNA-seq (snRNA-seq) has gained popularity since fresh and intact single cells are challenging to obtain in many applications. Hence, we tested the performance of SHERRY2 on snRNA-seq using single nuclei isolated from HEK293T cells. SHERRY2 detected 10,137 genes (RPM>1) on average at 1-million reads, which was 4,330 (74.6%) more than SmartSeq2, demonstrating that SHERRY2 had superior sensitivity for single nuclei (**Fig. 4A**). SHERRY2 still exhibited superior accuracy as it was significantly more correlated with NEBNext quantification results in comparison with SmartSeq2 (R=0.41 vs R=0.39) (**Fig. 4B**).

**Figure 4.**
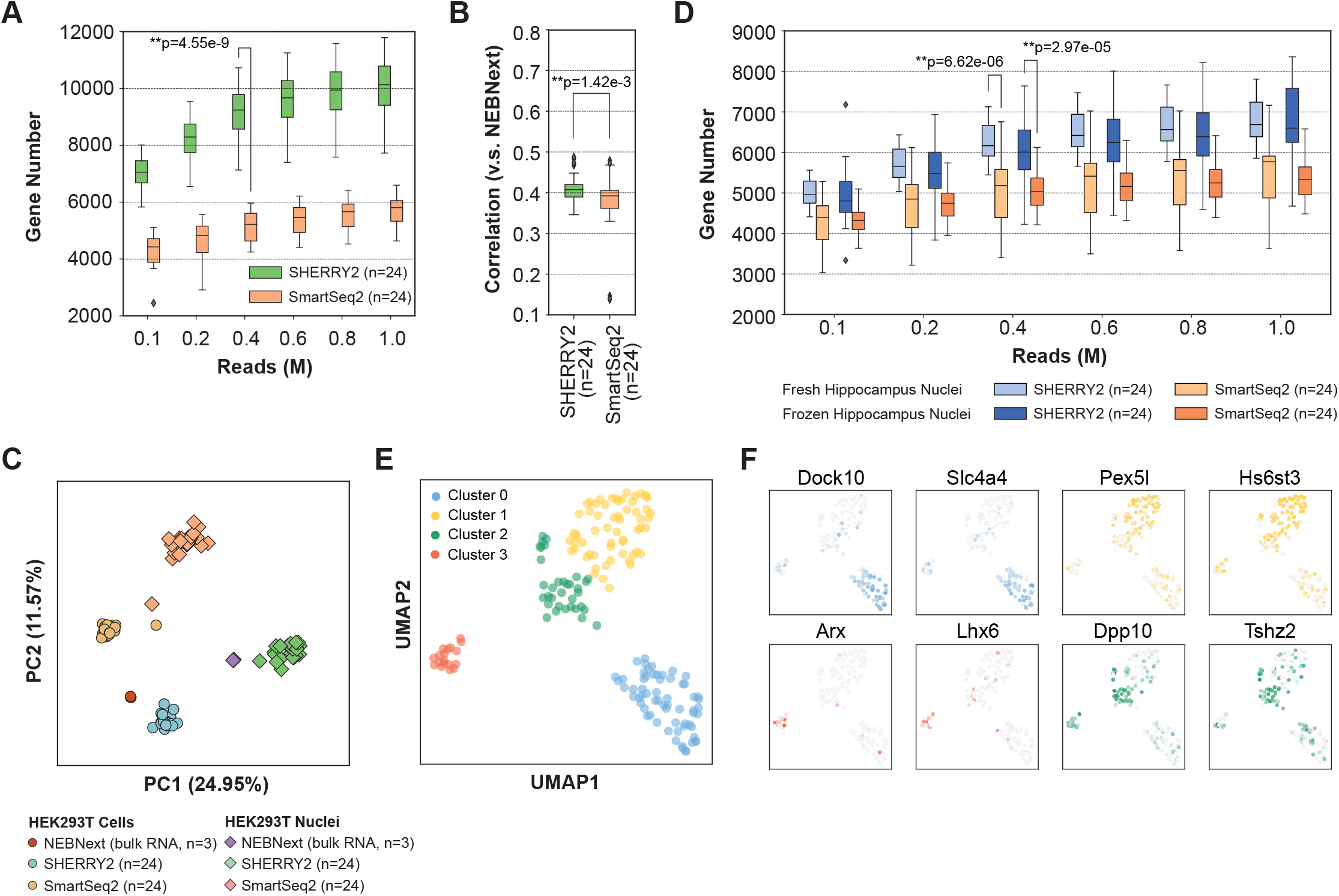
Sensitivity and accuracy of SHERRY2. (A) Gene number (RPM>1) of single HEK293T nuclei detected by SHERRY2 and SmartSeq2 when subsampling reads to 0.1,0.2, 0.4, 0.6, 0.8 and 1 million reads. (B) Gene expression correlation between single HEK293T nuclei and 200-ng RNA extracted from HEK293T nuclei. Single-nucleus data were acquired by SHERRY2 and SmartSeq2. Bulk RNA results were acquired by the standard NEBNext protocol. The correlation R-value was calculated by a linear fitting model with normalized gene counts. (C) Clustering of HEK293T cellular and nuclear RNA-seq data from SHERRY2, SmartSeq2 and NEBNext using principal component analysis. The analysis utilized differentially expressed genes (adjusted p-value < 1e-4 and fold change > 2) between cells and nuclei detected by NEBNext. (D) Gene number (RPM>1) of single neuron nuclei detected by SHERRY2 and SmartSeq2 when subsampling reads to 0.1, 0.2, 0.4, 0.6, 0.8 and 1 million reads. The nuclei were isolated from mouse hippocampi that were freshly prepared or previously frozen at −80°C. (E) Clustering of single hippocampal neuron nuclei visualized by UMAP plot. The snRNA-seq library was prepared by SHERRY2. The analysis utilized genes expressed (counts > 0) in more than 4 nuclei. (F) Marker gene expression of different cell types on UMAP plot from (E). The gradient colors correspond to the normalized counts of a specific gene ranging from 0 to 1. The p-values in (A, B, D) were calculated by the Mann-Whitney-U test.

The high accuracy and sensitivity of SHERRY2 allowed better distinction between HEK293T cells and their nuclei, which had minimal differences. We performed principal component analysis (PCA) using RNA-seq data from NEBNext, SHERRY2 and SmartSeq2 (**Fig. 4C**). Single cells and nuclei prepared by SHERRY2 were much closer in distance to the bulk RNA results in comparison with those prepared with SmartSeq2. In addition, the expression pattern of the differential genes identified by SHERRY2 was more similar to that of NEBNext in comparison with SmartSeq2 (**Additional file 1: Fig. S9**).

Furthermore, we compared the performance of these two methods with hippocampal neurons since snRNA-seq is a popular method for studies of brain tissue due to the technical challenge of isolating intact single neurons. We constructed snRNA-seq libraries of frozen and freshly prepared hippocampus with SHERRY2 and SmartSeq2. For both samples, SHERRY2 detected significantly more genes than SmartSeq2 (6,600 vs 5,331 at 1-million reads for frozen samples, 6,686 vs 5,769 at 1-million reads for fresh samples) (**Fig. 4D**). And still, Smart-seq2 tended to detect genes functionized in mitochondrion (**Additional file 1: Fig. S10A**). Next, we sequenced a small number of fresh hippocampal neurons (176 nuclei) (**Additional file 1: Fig. S10B**) with SHERRY2 and classified their cell types correctly. The nuclei were non-supervisedly clustered into 4 distinct groups (**Fig. 4E**), after which they were re-clustered using marker genes identified by sNuc-Seq [33] (**Additional file 1: Fig. S10C**). The two clustering results were highly consistent. Neurons within dentate gyrus (DG) and CA1, which occupy large areas of hippocampus, could be assigned to Cluster 0 and Cluster 1 respectively, according to the high expression of *Dock10, Slc4a4* and high expression of *Pex5l* and *Hs6st3* (**Fig. 4F**). However, CA3 pyramidal cells were not shown in our results, probably due to the small number of samples. Cluster 3 that were featured with enriched *Arx* and *Lhx6* could be annotated as GABAergic cells, which migrated from medial ganglionic eminence (MGE). Except for the forementioned markers, the expression patterns of these three clusters acquired from sNuc-seq and SHERRY2 were very similar (**Additional file 1: Fig. S10D**). Cluster 2 was found to consist of cells with relatively high expression of *Dpp10* and *Tshz2*, inferring that it might be contamination of cortex neurons. Moreover, our results revealed a long non-coding RNA (lncRNA) cluster [34] containing *Meg3, Rian (Meg8*) and *Mirg (Meg9*), which showed higher density in CA1 pyramidal cells and GABAergic cells while was relatively sparse in DG granule cells (**Additional file 1: Fig. S10E**).

## Discussion

SHERRY2 is a major improvement of our previously developed SHERRY [12], a Tn5 tansposase-based RNA-seq method that eliminates the second-strand complementary DNA synthesis. Although the original SHERRY protocol has shown satisfactory simplicity to construct RNA-seq libraries using low amount of starting material, the coverage bias at 3’-ends of transcripts and tagmentation-prone DNA contaminant make it challenging to work with single cells. In MINERVA [13], a direvative of SHERRY that specifically designed to work for metatranscriptome of COVID-19 clinical samples, we have explored the various conditions to reduce the DNA coverage. In SHERRY2, we further optimized the DNA reduction process and lead to a new protocol that can work for single cells and single nuclei, providing uniform coverage of whole transcirpts and resists to DNA contents.

There are three major advantages that SHERRY2 holds. First, SHERRY2 exhibits superior sensitivity and accuracy compared with SmartSeq2, a prevalent scRNA-seq method. What’s more, from sequencing data of single GC B cells and single neuron nuclei, we found that SmartSeq2 biasedly detected genes involed in mitochondrial components. Though more genes were obtained by SHERRY2, there was no specific functional enrichment of these genes (**Fig. 2G, Additional file 1: Fig. S10A**). Thus, SHERRY2 would have more chance to facilitate biological discoveries that relied on subtle changes. Recently SmartSeq3 [35], the upgraded protocol of SmartSeq2, has been reported to increase the gene detection sensitivity. We have also compared the scRNA-seq data of HEK293T cells produced by SHERRY2 and SmartSeq3. SHERRY2 is able to detect over 10,000 genes at around 1 million reads, while SmartSeq3 cannot aquire same number of genes even at 3-fold of sequencing depth (**Additional file 1: Fig. S11A**). Second, SHERRY2 retains great simplicity and expeditiousness, with the entire workflow taking around 3 hours and with all reactions performed in one tube. The swift experimental pipeline ensures less RNA degradation, eliminates the operational errors, and saves costs of supplies and labor. Third, SHERRY2 is highly robust and scalable. Procedural simplification not only reduces error cascade through step-wise operations, but also increases the tolerance of pipetting by offering easily-handled volumes, leading to a significantly higher repeatability when in comparison with SmartSeq3 (**Additional file 1: Fig. S11B**). Besides, SHERRY2 contains richer information about exon junctions and coding regions across full length transcripts, probably because SmartSeq3 is specifically optimized to quantify 5’-end of transcripts (**Additional file 1: Fig. S11C, S11D**).

SHERRY2 can be further developed to uncover more information from single cells. The simplicity and tolerance of protocol make it an ideal component to be incorporated into multi-omics studies. Moreover, since SHERRY2 actually contains the strand-specific information of the transcript since it builds libraries from RNA/DNA duplex directly. Therefore, SHERRY2 can be potentially modified to differentiate the transcriptional strand of DNA. In addition, barcoded Tn5 tagmentation [36, 37] may also be applied to SHERRY2 to realize assembling full-length RNA molecules. Interestingly, when examining reads generated by SHERRY2 and SmartSeq2, we find that the cleavage sites of Tn5 tend to exhibit different sequence bias on substrate DNA and RNA/DNA duplex, which might give hints to understand Tn5 mechanism (**Additional file 1: Fig. S12**).

There are a few remaining hitches of current SHERRY2 protocol that need to be fixed in the future. The slightly unsatisfactory mapping rate may be compensated by slightly more sequencing reads. Without cDNA enrichment, the exogenous DNA from environment or reagents still can be introduced after lysis step and easily tagged by Tn5, thus impairing the performance of RNA-seq of low RNA content single cells or nuclei. Besides, it’s still challenging to capture the complete 5’-end regions of transcripts for the limited processivity of reverse transcriptase. For example, CDR1 and CDR2 sequence in Igh variable regions cannot be acquired for all GC cells (**Additional file 1: Fig. S8C**).

## Conclusions

We present SHERRY2, an RNA-seq method designed for single cells and single nuclei. SHERRY2 is based on the direct tagmentation function of Tn5 transposase for RNA/DNA hetero-duplexes, and overthrows prevalent single cell RNA-seq chemistries which typically require pre-amplification of full-length transcripts, thus greatly improving the sensitivity of gene detection and eliminating the sequence-dependent bias. As a result, SHERRY2 can reveal expression dynamics of transcription factors and lncRNAs, both of which typically harbor essential biological functions while at low abundance. Meanwhile, SHERRY2 maintains the simplicity of operation, with whole process completed in one pot within 3 hours, and hence elevates the throughput to a few thousand single cells/nuclei per experimental batch. As the simplest protocol of large-depth scRNA-seq, SHERRY2 has been validated in various challenging samples, and can be seamlessly integrated into wide range of applications.

## Methods

### Cell culture

HEK293T cell line was purchased from ATCC and incubated at 37°C with 5% CO_2_ in Dulbecco’s Modified Eagle Medium (DMEM) (Gibco, 11965092), which was supplemented with 10% fetal bovine serum (FBS) (Gibco, 1600044) and 1% penicillin-streptomycin (Gibco, 15140122). Cells were dissociated by 0.05% Trypsin-EDTA (Gibco, 25300062) at 37°C for 4min and washed by DPBS (Gibco, 14190136).

For DNA or RNA extractions, we took ~10^6^ suspended cells, and followed the guideline of PureLink Genomic DNA Mini Kit (Invitrogen, K182002) or RNeasy Mini Kit (Qiagen, 74104). The extracted RNA was further dealt with 20U DNase I (NEB, M0303) for removal of DNA and re-purified by RNA Clean & Concentrator-5 kit (Zymo Research, R1015).

For single nuclei preparation, we followed the guideline of Nuclei EZ Prep kit (Sigma, NUC-101) and resuspended the nuclei into DPBS. Both single cells and single nuclei were sorted by FACSAria SORP flow cytometer (BD Biosciences).

### Mice

For samples of germinal center B cells, C57BL/6 mice were originally from the Jackson Laboratory. 6-12 week-old, age- and sex-matched mice were used for the experiments.

For samples of hippocampus nuclei and lymphocytes, aged (2-months old) male C57BL/6 mice were used and obtained from Charles River Laboratories.

All mice were maintained under specific pathogen-free conditions and used in accordance of governmental, Tsinghua University and Capital Medical University guidelines for animal welfare.

### GC light zone B cells preparation and sorting

To generate T-cell dependent GC responses in B6 mice, 100μg NP-KLH (Biosearch Technologies, N-5060-5) plus 1μg LPS (Sigma, L6143) emulsified in 100μl 50% alum (Thermo, 77161) were utilized for intraperitoneal immunization.

Spleens isolated from 4 mice of 13-days post immunization were placed on a 70μm cell strainer (Falcon, 08-771-2), which was previously wetted with MACS buffer (1% FBS and 5mM EDTA in PBS), and minced by flat end of the plunger of 2ml syringes (Becton Dickinson, 301940). The splenocytes passed through the strainer with MACS buffer into a 50ml-tube. The mixed red blood cells were then lysed by ACK lysis buffer (Thermo, A1049201). The cell suspension was further incubated with biotinylated 4-Hydroxy-3-iodo-5-nitrophenylacetyl (NIP)15-BSA (Biosearch Technologies, N-1027-5) for 1.5h, and enriched by Anti-biotin cell isolation kit (Miltenyi Biotec, 130-090-485) to get NP-reactive cells.

The enriched cells were blocked with 20μg/ml 2.4G2 antibody (BioXCell, BE0307) and subsequently stained with APC-Cy7 (anti-B220, BD Biosciences, 552094), PE-Cy7 (anti-CD95, BD Biosciences, 557653), eF450 (anti-GL7, eBioscience, 48-5902-82), APC (anti-CD86, eBioscience, 17-0862-82) and PE (anti-CXCR4, BioLegend, 146505). Also, 7-AAD (Biotium, 40037) was stained to exclude dead cells. All staining reactions were incubated in MACS staining buffer (1% FBS and 5mM EDTA in PBS) for 30min on ice, followed by 3 times of washings. As gated in **Additional file 1: Fig. S5B**, single GC Light Zone B cells (B220^+^ GL7^+^ Fas^+^ CD86^+^ CXCR4^-^) were sorted into in lysis buffer using Aria III flow cytometer (BD Biosciences).

### Lymphocyte cells preparation and sorting

The retro-orbital blood was taken from the eyeball of ether-anesthetized mice and dipped into K2EDTA tube (BD Vacutainer, 367525). PBS was added to dilute blood at ~50%. 1ml diluted blood was transferred into a clean 15ml-tube and incubated with 9ml 1x red blood cells lysing solution (BD Pharm Lyse, 555899) at room temperature for 15min avoiding light. The resulted cell suspension was washed twice by PBS containing 1% BSA at 200g for 5min, followed by staining with SYTOX green (Thermo, S7020) to identify intact cells. Single lymphocytes were sorted with FACSAria SORP flow cytometer according to the gates shown in **Additional file 1: Fig. S5A**.

### Hippocampal nuclei preparation and sorting

The isolated hippocampus tissue was transferred into a Dounce homogenizer (Sigma, D8938) containing 2ml of EZ Lysis Buffer (Sigma, NUC-101). The tissue was carefully dounced for 22 times with pestle A followed by 22 times with pestle B, then transferred to a 15ml-tube. Next, 1ml of EZ lysis buffer was added into the Dounce homogenizer to resuspend residual nuclei, then transferred to the same 15ml tube. The samples were centrifuged at 300g for 5 min. Supernatant was removed and the pellet was resuspended in 100μl of ice-cold PBS (Gibco, 10010023) with 1% BSA (NEB, B9000S) and 20U RRI (Takara, 2313). 40μm FlowMi cell strainers were firstly wetted with PBS and filtered the resuspended nuclei into 1.5 ml Eppendorf tubes. The nuclei were further washed by PBS (1% BSA).

To enrich neuron nuclei, 1,000-fold diluted mouse Anti-NeuN antibody (Millipore, MAB377) was added into 0.5ml nuclei suspension and incubated with the nuclei at 4°C for 30min. The nuclei were then stained with 1000-fold diluted Goat anti-Mouse IgG (H&L) antibody (Abcam, ab150113) and washed with PBS (1% BSA). The whole process was on ice. As gated in **Fig. S10B**, single neuron nuclei were sorted with FACSAria SORP flow cytometer.

For frozen samples, hippocampus tissues were previously flash frozen in liquid nitrogen, and stored in −80°C. Before single nuclei preparation, they were thawed on ice totally.

### DNA carrier preparation

100-ng pTXB1 plasmids were firstly linearized by 10U XbaI (NEB, R0145S) at 37°C for 1h and purified by Zymo columns. Then we took 0.5-ng linearized plasmids for multiple displacement amplification (MDA), with all dTTPs replaced by dUTPs. Specifically, the 1μl DNA was mixed with 22μl reaction buffer containing 1x phi29 reaction buffer (NEB, M0269S), 20μM random primers (Thermo, SO181) and 1mM dNTP (NEB, N0446S and N0459S), then they were incubated at 98°C for 3min and immediately cooled down at 4°C for 20min. 2μl phi29 DNA polymerase was added and the amplification was carried out at 30°C for 5h, terminated at 65°C for 10min. The products were purified by Zymo columns.

### Generation of RNA-seq library

We constructed NEBNext libraries with 200- and 10-ng RNA by following the guideline of NEBNext Ultra II RNA Library Prep Kit for Illumina kit (NEB, E7770). SmartSeq2 libraries with single cells were prepared following the protocol that was reported by Picelli, S. *et al* [3]. 10X libraries of 10,000 single hippocampal nuclei were constructed by Chromium Single Cell 3’ Reagent Kits (v3.1).

For scRNA-seq library of SHERRY2, single cells were sorted into 96-well plates containing 2μl lysis buffer which consisted of 0.5% Triton X-100 (Sigma, T9284), 2U SUPERaseIn RNase Inhibitor (Thermo, AM2694), 0.2U AG DNase I (Thermo, 18068015). The plates were immediately spun down and incubated at 20°C for 10min for DNA digestion. The plates could be stored at −80°C or proceeded with next step. 2μl inactivation buffer containing 5μM OligodTs (T30VN, Sangon), 5mM dNTPs and 1mM EDTA (Thermo, AM9260G) was then added and the reaction was incubated at 65°C for 10min and 72°C for 3min to facilitate RNA denaturation at the same time. Next, RT was performed by adding 6μl RT mix (70U SuperScript IV (Thermo, 18090050), 1.7x SSIV buffer, 8.3mM DTT, 10U RRI, 1.7M Betaine (Sigma, B0300)) and incubated at 50°C for 50min, then inactivated the reverse transcriptase at 80°C for 10min. The resulted RNA/DNA hybrids mixed with 10-pg DNA carriers were tagmented by 0.05μl TTE Mix V50 (Vazyme, TD501) at 55°C for 30 min, through adding 10μl reaction mix containing 2x TD buffer (20mM Tris-HCl (ROCKLAND, MB-003), 10mM MgCl2 (Thermo, AM9530G), 20% N,N-Dimethylformamide (Sigma, D4551)), 16% PEG8000 (VWR Life Science, 97061), 0.5mM ATP (NEB, P0756), 8U RRI. 6U Bst 3.0 DNA polymerase (NEB, M0374M) within 1 x Q5 high-fidelity master mix were utilized to repair the gap left by V50 at 72°C for 15min, followed by 80°C for 5min to terminate the reaction. Finally, 3μl indexed primers mix (Vazyme, TD203) and 3μl Q5 mix were added to perform PCR amplification. PCR cycled as following: 98°C 30s for initial denaturation, 21 cycles of 20s at 98°C, 20s at 60°C and 2min at 72°C, 72°C 5min for final extension. The indexed products were merged and purified at 0.75x with VAHTS DNA Clean Beads (Vazyme, N411).

Libraries were quantified with Qubit 2.0 and their fragment length distributions were checked by Fragment Analyzer Automated CE System. Libraries were sequenced by Illumina NextSeq 500 or NovaSeq S4.

### RNA-seq data analysis

#### Data quality

Adaptors, poly(A/T) sequences were trimmed, bases with quality less than 20 and reads shorter than 20 bases were removed from the raw sequencing data by Cutadapt (v1.15) [38]. Clean reads were mapped to indexed genome (human: Gencode.v31, mouse: Gencode.vM23) by STAR (2.7.1a) [39]. Only unique alignment was utilized for downstream analysis. The mitochondrial and ribosomal ratios were counted with samtools (v1.10) [40]. The ratios of coding region, UTR, intron and intergenic region were counted with Picard tools (v2.17.6). Exonic rate was defined as sum of coding region and UTR ratios. For cells, Cufflinks (v2.2.1) [41] with exon annotations of protein coding genes were used to count gene number (FPKM>1). For nuclei, genes (RPM>1) were counted by featureCounts (v1.5.1) [42] with transcript annotations. Coverage across gene body was calculated by RSeQC (v.2.6.4) [43]. The coverage uniformity was integral area between coverage curve and x-axis normalized by 100.

#### Gene ontology analysis

We used genes that were detected in cell A while missed by cell B as “study”, and combined the “study” genes with genes detected by cell B as “backgroud”. The gene ontology analysis was performed by GOATOOLS (v1.2.3) [44] and repeated between every two cells from different methods. GO terms (excluding electronic annotations) with adjusted p-value less than 0.01 were counted. All cells were firstly downsampled to 500K or 1M total reads.

#### Clustering and marker genes

For scRNA-seq and snRNA-seq, clustering followed the basic tutorials of Scanpy (v1.8.1) [45]. The cell type annotations were through manually checking expression of well-known marker genes. Marker genes that identified by SHERRY2 should satisfy following conditions: 1) adjusted p-values calculated by Mann-Whitney-U test were less than 1e-3; 2) foldchanges were greater than 1.5 or less than 0.67; 3) The average normalized counts of up-regulated gene in the cell type, or down-regulated gene in the rest of cell types was greater than 0.3. For NEBNext, DESeq2 (v1.22.2) [46] was utilized to identify the differentially expressed genes (adjusted p-value < 1e-4, foldchange > 2).

#### RNA Velocity

Splicing and unsplicing mRNA were quantified by Velocyto (v0.17.17) [10] with unique alignment. The generated loom file was utilized by scVelo (v0.2.4) [47] to profile velocity dynamics based on clustering results of Scanpy.

#### BCR assembly

BCR sequences of each cell was assembled by MIXCR (v3.0.13) [48] with clean reads. The assembled BCR were realigned by IgBlast (v1.17.1) [49] to determine clone types.

#### Exon usage

The frequency of exon usage in each cell was calculated by BRIE (v2.0.5) [50]. For each exon, cells satisfying following conditions were retained: 1) counts of gene which included the exon were greater than 10; 2) exon regions sided by the specific exon should be covered by greater than 50% with uniquely aligned reads; 3) at least one read should detect junctions involved in this exon splicing events. Pairwise comparison of exon usage frequency was made between cell types which contained greater than 10 cells using Mann-Whitney-U test. The exons with p-value less than 0.05 was further checked in IGV viewer to check whether transcript coverage was consistent with usage frequency. The passed ones were considered as significantly biased among cell types.

#### SmartSeq3 data reanalysis

SmartSeq3 [35] sequencing data of 117 single HEK293T cells was downloaded from ArrayExpress. The UMI and tag sequences at 5’-end were firstly removed. Merged 5’-end reads and internal reads were then analyzed using pipeline described in Data quality.

## Declarations

### Consent for publication

Not applicable

### Availability of data and materials

All data generated or analysed during this study are included in this published article, its supplementary information files and publicly available repositories. The sequence data reported in this study have been deposited in the NCBI Sequence Read Archive (assession no. PRJNA879104).

### Competing interests

The authors declare that they have no competing interests.

### Funding

This work was supported by the National Key Research and Development Program of China (2018YFA0108100 to Y.H.); the National Natural Science Foundation of China (22050002, 21927802 to Y.H.); the Beijing Municipal Science and Technology Commission (Z201100005320016 to Y.H.); and the Shenzhen Bay Laboratory. Funding for open access charge: Beijing Municipal Science and Technology Commission.

### Authors’ contributions

Y.H. and J.W. conceived the study; L.D., B.L., Y.L., S.Z., Y.P., and J.S. performed exepriemtns; L.D. performed data analyses; C.Z. and J.S. provided samples; L.D., B.L., J.W., H.Q., J.S., and Y.H. wrote manuscript with input from all authors; Y.H., J.W., and H.Q. supervised all aspects of this study.

## Acknowledgements

We thank Chenyang Geng and Yan Chen from the Peking University High-throughput Sequencing Center and Biomedical Pioneering Innovation Center for experimental assistance. This work was supported by the National Key Research and Development Program of China (2018YFA0108100 to Y.H.), National Natural Science Foundation of China (22050002, 21927802 to Y.H.), Beijing Municipal Science and Technology Commission (Z201100005320016 to Y.H.), Beijing Advanced Innovation Center for Genomics, and Shenzhen Bay Laboratory.

## Notes

### Competing Interest Statement

The authors have declared no competing interest.

### Summary of Updates

Title updated; Figures updated; Supplemental files updated.

