## Additional file 1 for "Rapid and sensitive single cell RNA sequencing with SHERRY2"

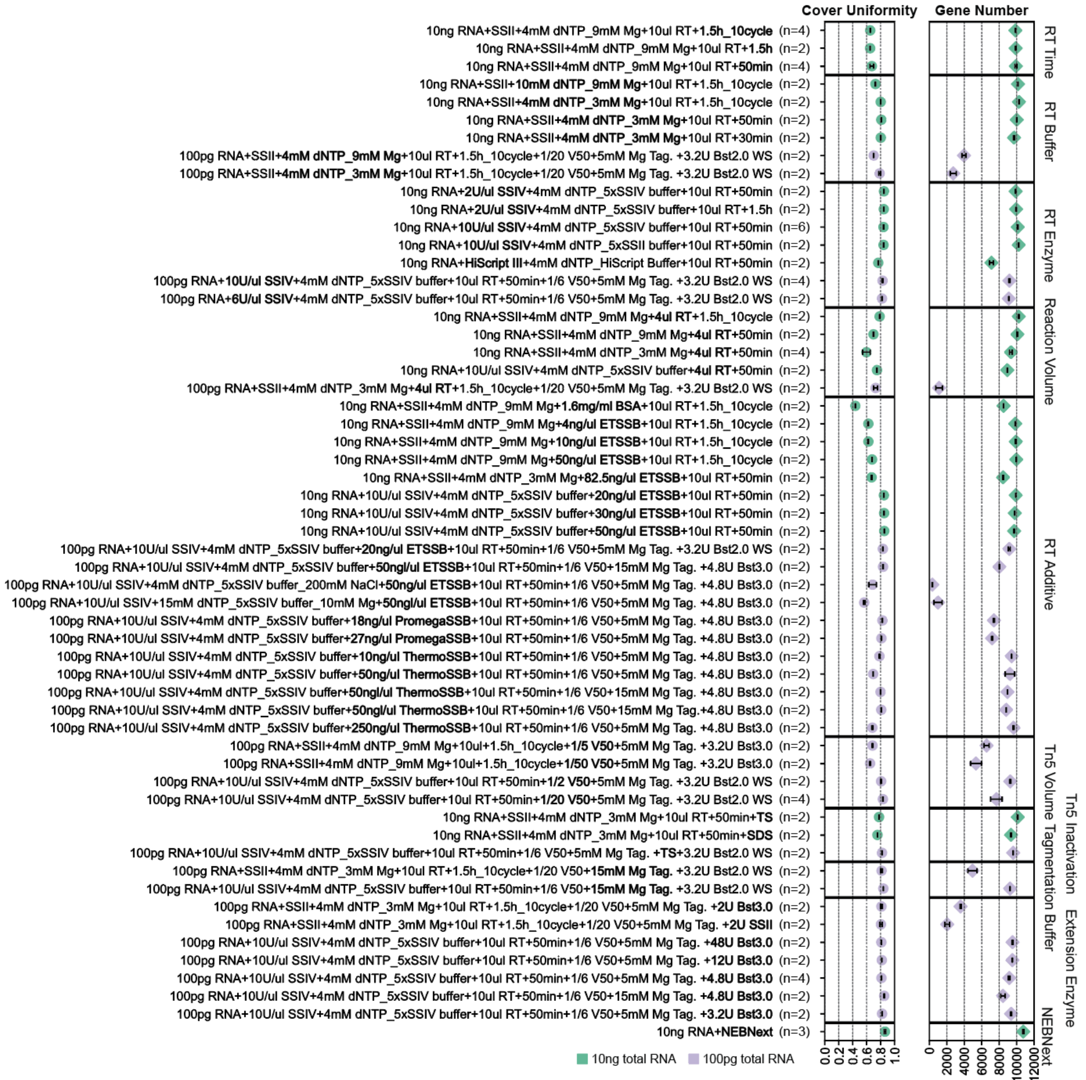


**Fig. S1.** Conditions for bulk RNA. RNA-seq library quality of 10-ng (green) and 100-pg (purple) HEK293T RNA samples when testing different conditions of the SHERRY protocol. The coverage uniformity across gene body and gene number (FPKM>1) at 0.25-million reads are shown here. The bolded fonts of the labels indicate key variations compared to the basic protocol. The NEBNext standard protocol was performed with 10-ng HEK293T RNA as a reference. The error bars show the standard deviation.

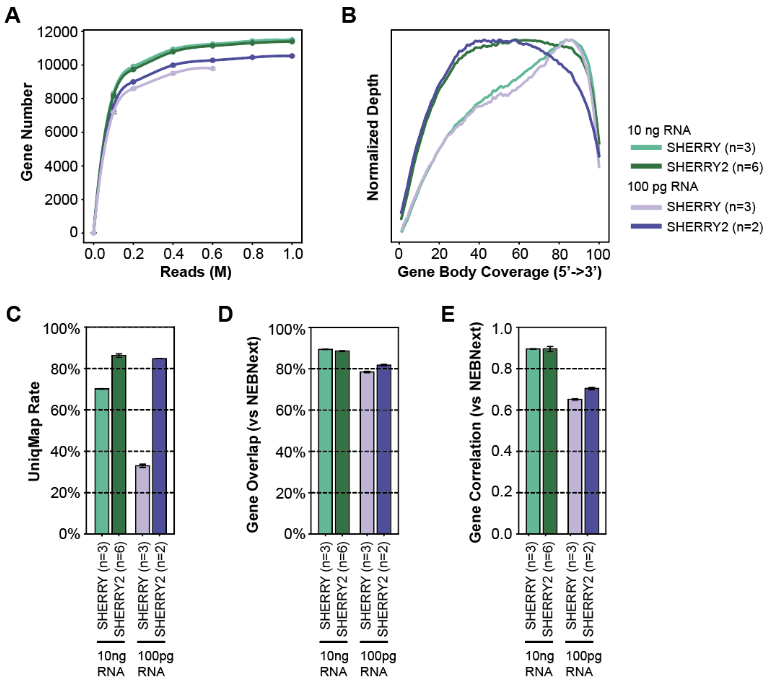


**Fig. S2.** Improved performance of SHERRY2 on bulk RNA. (A) Saturation curve of SHERRY and SHERRY2. The gene number (FPKM>1) is shown for sequencing at 0.2, 0.4, 0.6, 0.8, and 1.0 million reads. (B) Gene body coverage detected by SHERRY and SHERRY2. (C) Ratio of reads that uniquely mapped to the human genome. (D) Ratio of expressed genes (FPKM>1) in the NEBNext results that overlapped the SHERRY and SHERRY2 results. (E) Correlation of overlapped gene counts between NEBNext and the two protocols: SHERRY and SHERRY2. The inputs of SHERRY and SHERRY2 in (A-E) were 10-ng and 100-pg HEK293T RNA. The input of NEBNext in (D, E) was 200-ng HEK293T RNA. The error bars in (C-E) show the standard deviation.

A

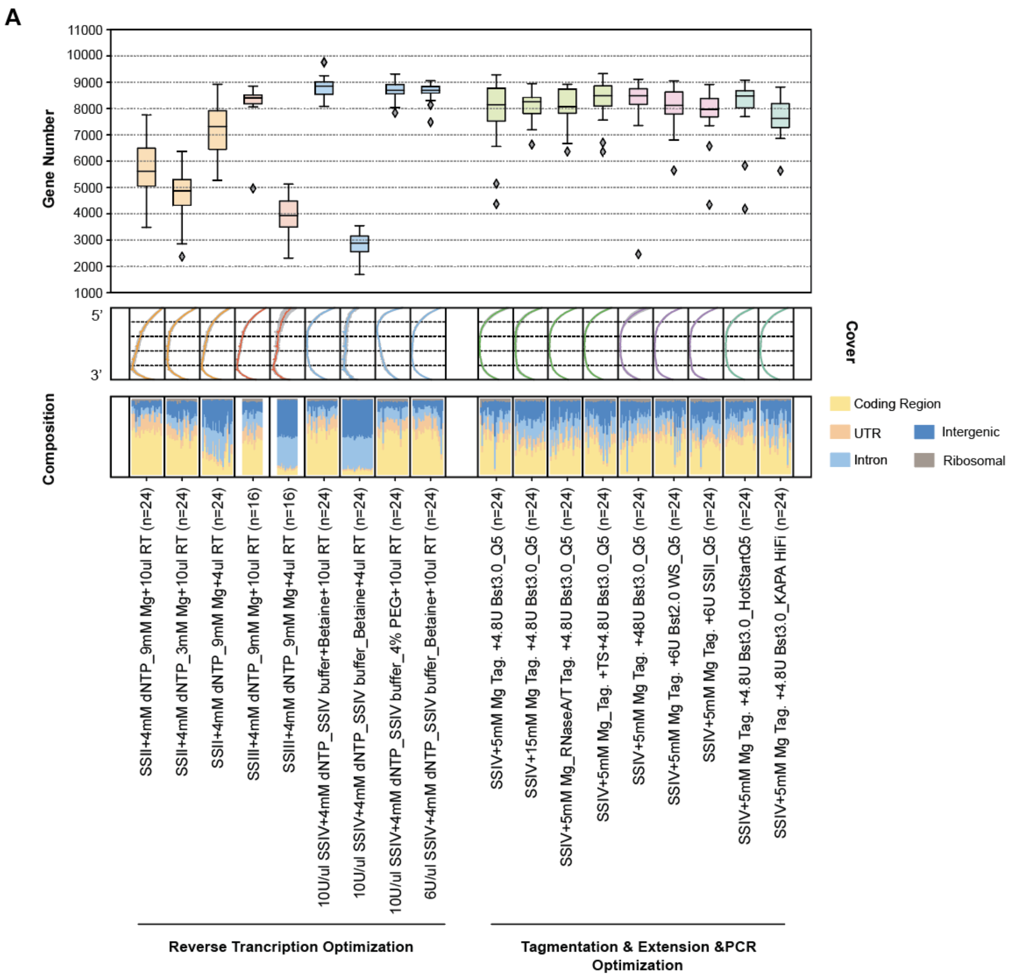


B

| | SHERRY | SHERRY2 | SmartSeq2 |
|---|---|---|---|
| **Lysis+RT** | ~$3.94 | ~$5.27 | ~$4.04 |
| **Pre-amplification** | - | - | ~$2.09 |
| **Library Prep** | ~$6.46 | ~$3.41 | ~$42.10 |
| **Total** | ~$10.40 | ~$8.68 | ~$48.23 |

**Fig. S3.** Conditions for single cells. (A) scRNA-seq library quality of single HEK293T cells when testing different conditions of SHERRY protocol. The gene number (FPKM>1) at 0.25-million reads, gene body coverage and composition of reads mapped to different genome regions are shown. The colors of the boxplot and coverage plot indicate key variations in different steps. The gray region of the coverage plot shows the standard deviation of normalized depth among replicates. (B) Costs for library preparation when using SHERRY, SHERRY2 and SmartSeq2.

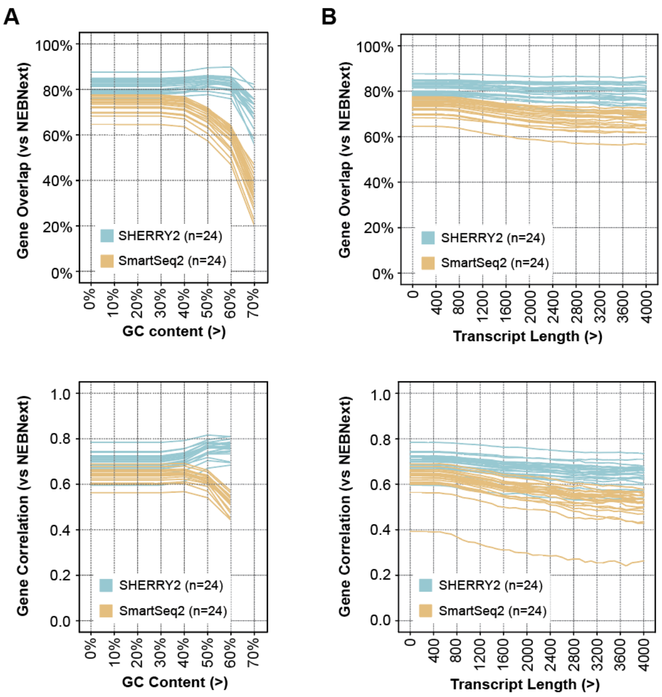


**Fig. S4.** Accuracy of SHERRY2 for single cells. (A) Ratio of expressed genes (FPKM>1) with different GC-content in the NEBNext results that overlapped by SHERRY2 and SmartSeq2, and the correlation of overlapped gene expression results between NEBNext and the two scRNA-seq methods. The x-axis indicates the minimum level of GC-content. Fewer than 100 genes had GC content >70%, and they were not included in the analysis. Each line represents one cell. (B) Gene overlap ratios and correlation of gene counts between NEBNext and the two scRNA-seq methods, utilizing genes with different transcript lengths from the NEBNext results. The inputs of SHERRY2 and SmartSeq2 were single HEK293T cells. The input of NEBNext was 200-ng HEK293T RNA. The correlation R-value was calculated by a linear fitting model.

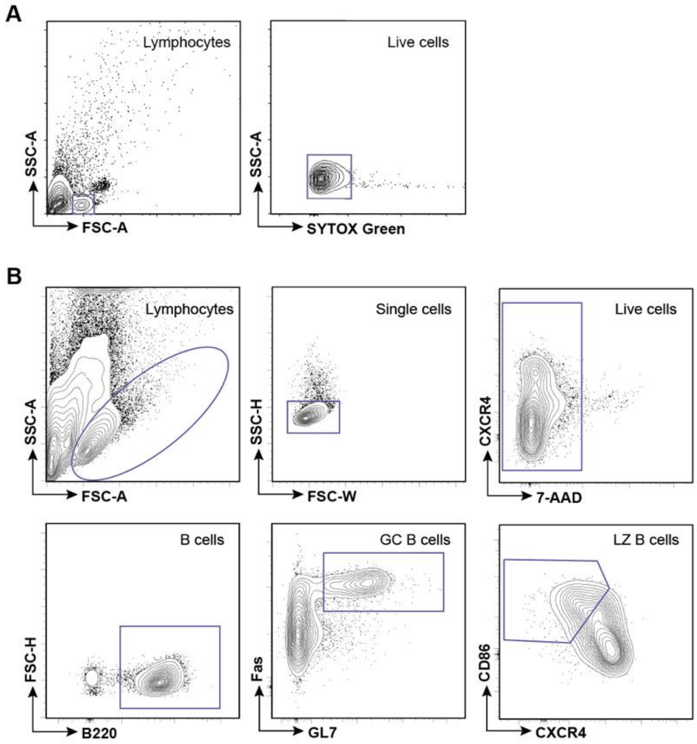


**Fig. S5.** Flow cytometry gating of single lymphocytes and single GC B cells. Flow cytometry gating of single lymphocyte cells that were isolated from murine eyeball blood (A) and single B cells from murine GC light zones (B).

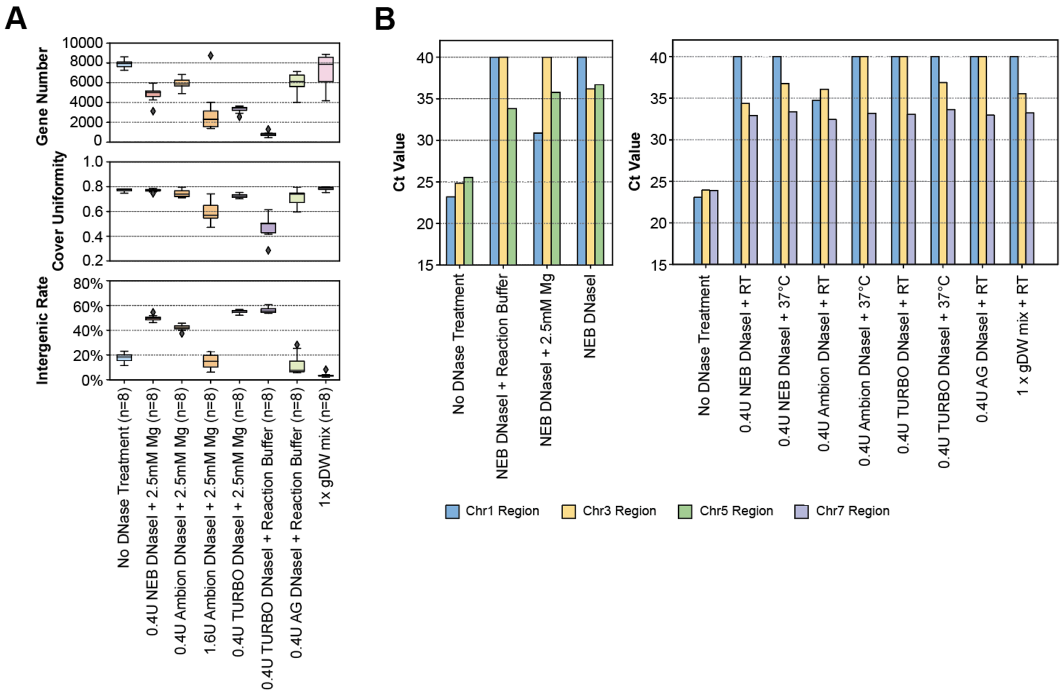


**Fig. S6.** DNase activity and performance in SHERRY2 library construction. (A) Library quality of SHERRY2 tested with single HEK293T cells and different DNases. The gene number (FPKM>1) at 0.25-million reads, gene body coverage uniformity and percentage of reads that were mapped to intergenic regions are shown. The x-axis labels indicate the amount of each DNase and its reaction buffer. (B) Left panel: qPCR quantification of 15-ng digested HEK293T genomic DNA by 0.4U NEB DNase I with the recommended reaction buffer, either containing 2.5 mM $Mg^{2+}$ or without any divalent ions. The digested products were split into three parts, and ct values were measured using primers designed for the chr1, chr3 or chr5 region. Right panel: qPCR quantification of 30-ng digested HEK293T genomic DNA. The x-axis labels indicate the amounts of each DNase and the reaction temperature (RT indicates room temperature). The digestions were all performed without any added divalent ions. The digested products were split into three parts, and ct values were measured using primers designed for the chr1, chr3 or chr7 region.

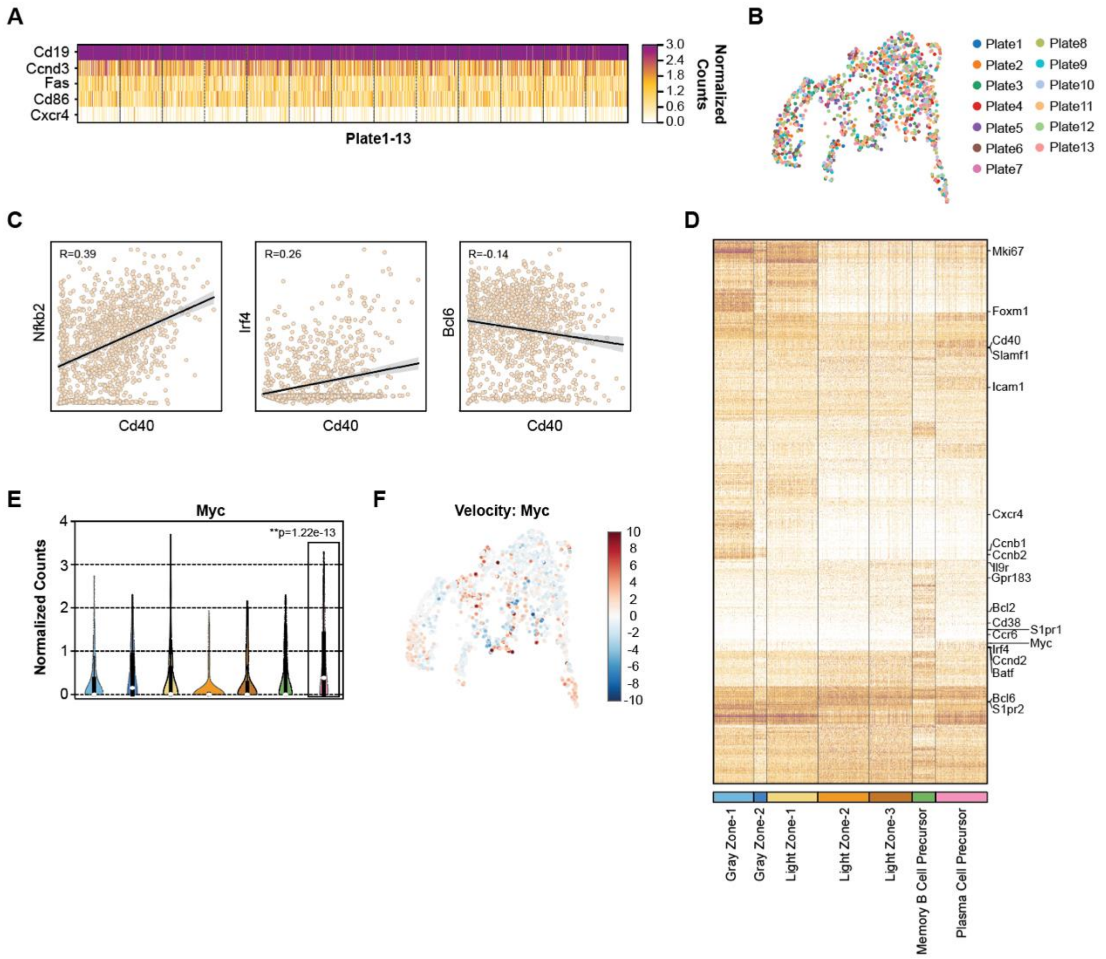


**Fig. S7.** GC information gained from SHERRY2. (A) Heatmap of the counts of the gating genes shown in Fig. S5B, which were used to isolate single B cells from murine GC light zones. (B) Single cells sorted in different 96-well plates were labeled with different colors on a UMAP plot showing clusters from the GC light zone in Fig. 3A. (C) Expression of Cd40 correlated with expression of *Nfkb2, Irf4* and *Bcl6*. R-value was calculated by a linear fitting model with normalized gene counts. (D) Up- or down-regulated genes (adjusted p-value < 1e-3, fold change > 1.5 or <0.67) identified by SHERRY2 across GC light zone cell types. The labeled genes were previously reported for the cell types. (E) Violin plot of gene *Myc* counts within each GC light zone cell type. The colors correspond to the cell types annotated in Fig. 3A. The p-values between plasma cell precursors and other cell types were calculated by the Mann-Whitney-U test. (F) RNA velocity of the *Myc* gene projected on a UMAP plot.

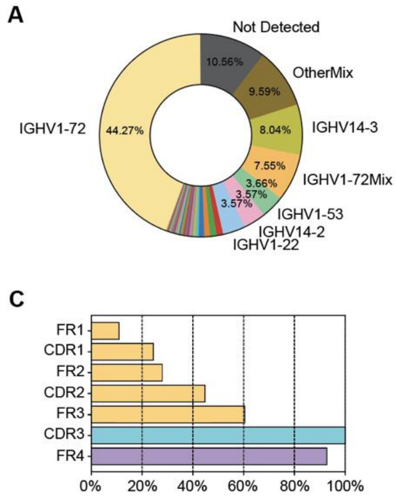


**Fig. S8.** BCR sequences of single GC B cells identified by SHERRY2. (A) Usage frequency of IgH variable genes that were assembled from the reads of each single GC light zone B cell. "Mix" indicates that more than 1 IgH variable gene was assigned, or more than 1 heavy chain sequence was assembled and also assigned to different genes, with a frequency less than 80%. "IGHV1-72Mix" indicates that one of the assigned genes was IGHV1-72. (B) Usage frequency of light chain types that paired with IGHV1-72 sequences. (C) Proportions of cells that covered different regions within IgH variable genes among cells for which IgH sequences could be assembled.

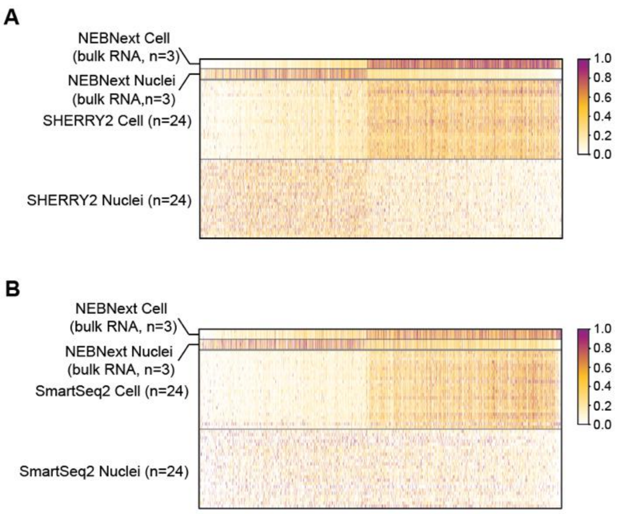


**Fig. S9.** Expression pattern of differential genes between cells and nuclei in SHERRY2. Heatmap of the counts of differentially expressed genes between HEK293T cells and nuclei in the NEBNext, SHERRY2 (A) and SmartSeq2 (B) results. The differentially expressed genes were identified by the NEBNext method with 200-ng RNA input and sorted by log2 fold-change. The inputs of SHERRY2 and SmartSeq2 were single HEK293T cells and nuclei. Each line represents a replicate.

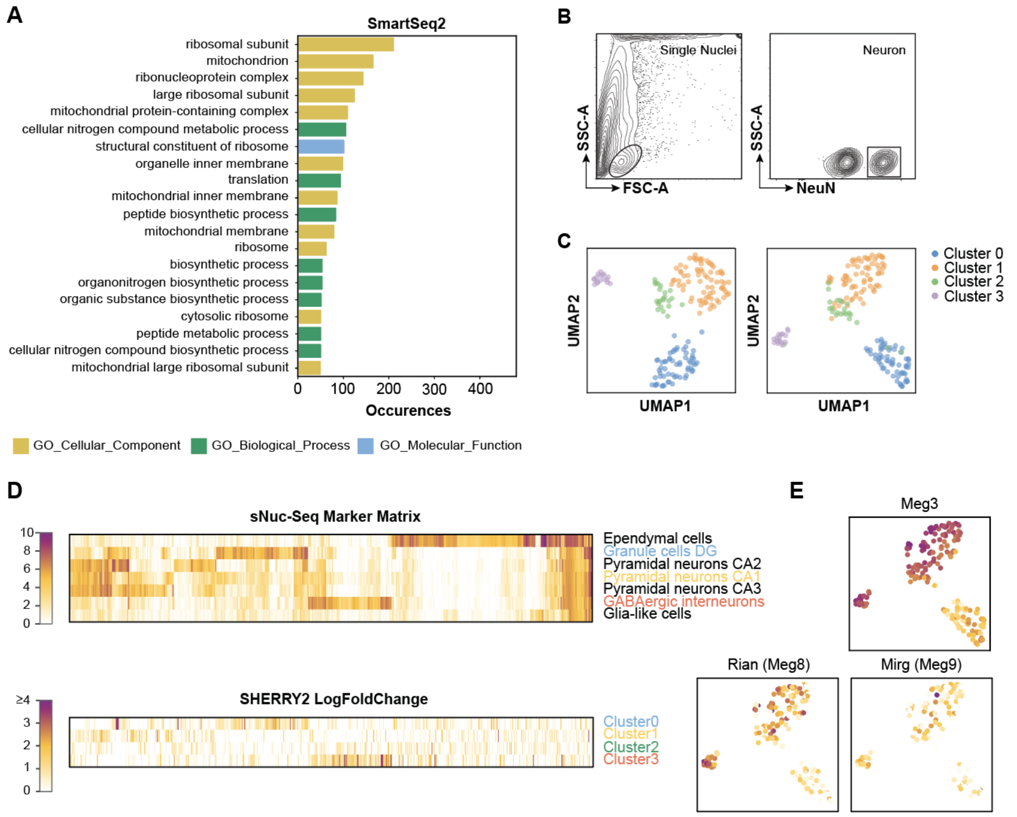


**Fig. S10.** Clustering and annotation of hippocampal nuclei. (A) Gene ontology analysis of genes that detected by SmartSeq2 while missed by SHERRY2 for single freshly-prepared hippocampal nuclei. The top 20 most commonly occurred GO terms were shown. (B) Flow cytometry gating of single neuron nuclei that were isolated from mouse hippocampus. C) The left panel shows a UMAP projection of the neuron nuclei clustered using marker genes of hippocampal cell types reported by Habib, N. *et al* (16). The right panel shows the nuclei positions in Fig. 4E, and each nucleus is colored using the same labeling scheme used in the left panel. The snRNA-seq libraries were prepared by SHERRY2. (D) Heatmaps of hippocampal maker gene counts acquired from Habib, N. *et al* (16) and their log fold change across clusters that were identified by SHERRY2. (E) Relative counts (normalized in range of 0-1) of 3 lncRNA projected on the UMAP plot of Fig. 4E.

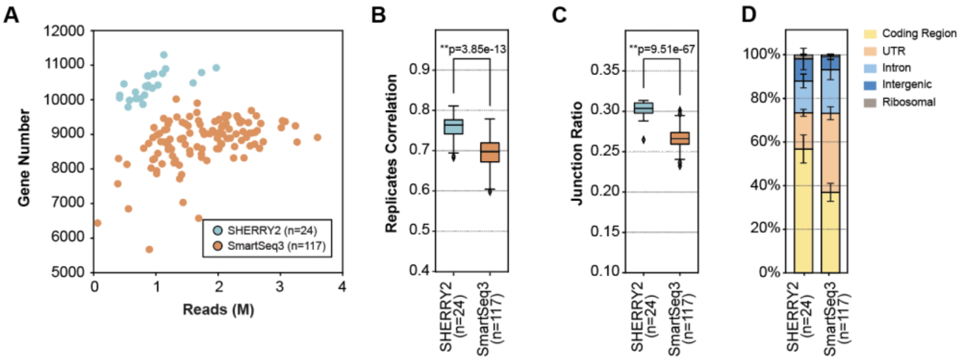


**Fig. S11.** Comparison of SHERRY2 and SmartSeq3. (A) Gene number (RPKM>1) of single HEK293T cells detected by SHERRY2 and SmartSeq3 with different sequencing depth. (B) Pairwise correlation of gene expression within replicates. The correlation R-value was calculated by a linear fitting model with normalized counts of overlapped genes. (C) Percentage of exonic reads that contained junctions. (D) Components of reads that were mapped to different regions of the genome. The p-values in (B, C) were calculated by the Mann-Whitney-U test.

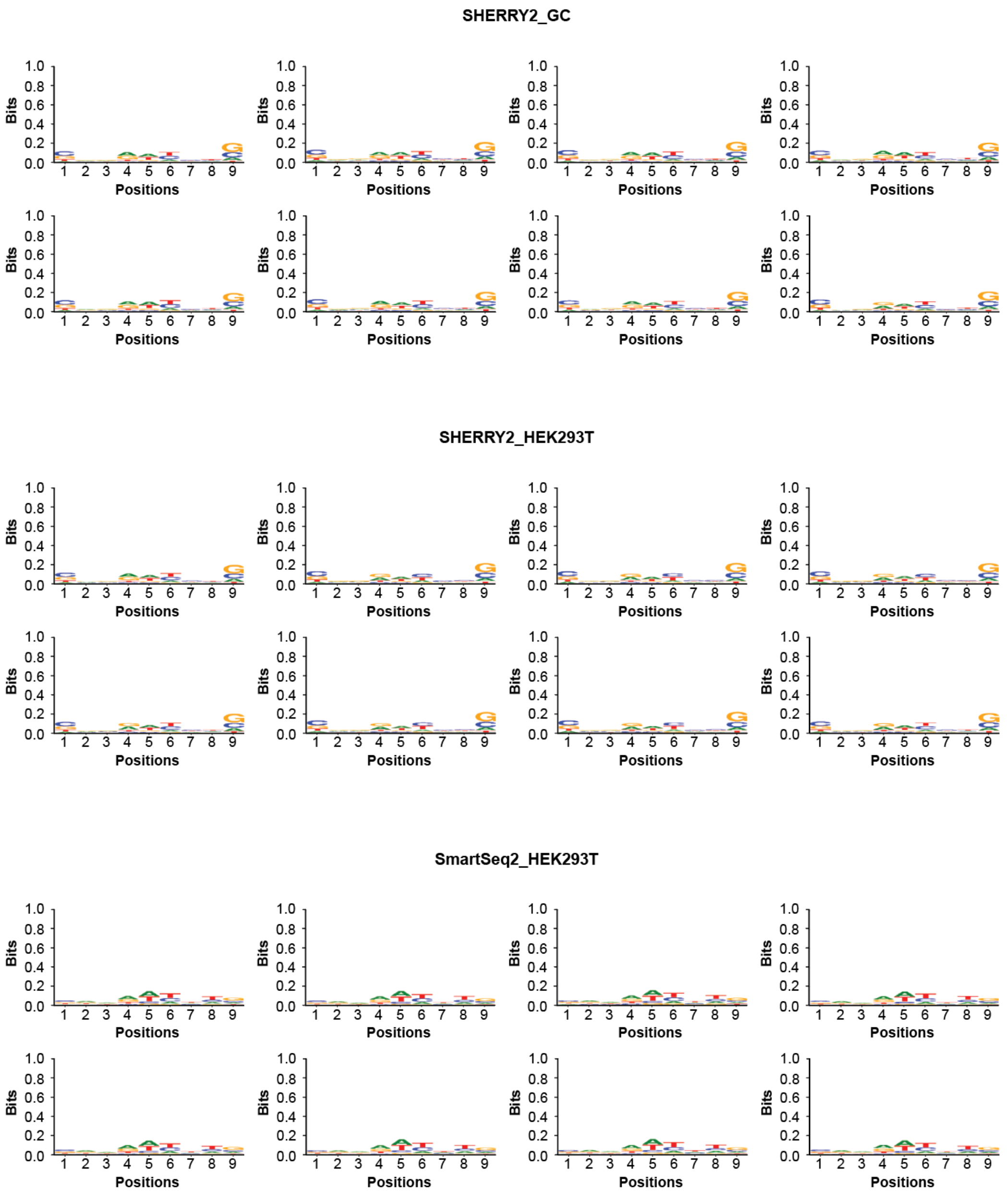


**Fig. S12.** Sequence bias in tagmentation of different substrates. Motif of first 9 bp within Read1 that were mapped to exonic regions. The first two panels showed motifs in SHERRY2 results that sequenced single murine GC B cells and HEK293T cells. The last panel showed motifs in SmartSeq2 results that sequenced single HEK293T cells.
